# Hydrogen isotope composition of *Thermoanaerobacterium saccharolyticum* lipids: comparing wild type to a *nfn-* transhydrogenase mutant

**DOI:** 10.1101/138651

**Authors:** William D. Leavitt, Sean Jean-Loup Murphy, Lee R. Lynd, Alexander S. Bradley

**Affiliations:** Department of Earth and Planetary Sciences, Washington University in St. Louis, Saint Louis, MO 63130 USA; Department of Earth Sciences, Dartmouth College, Hanover, NH 03755 USA; Department of Biological Sciences, Dartmouth College, Hanover, NH 03755 USA; Thayer School of Engineering, Dartmouth College, Hanover, NH 03755 USA; BioEnergy Science Center, Oak Ridge National Laboratory, Oak Ridge TN 37830, USA; Division of Biology and Biomedical Sciences, Washington University in St. Louis, Saint Louis, MO 63130 USA

**Author notes:** Correspondence authors e-mail addresses (W.D. Leavitt, (A.S. Bradley).

## Abstract

The ^2^H/^1^H ratio in microbial fatty acids can record information about the energy metabolism of microbes and about the isotopic composition of environmental water. However, the mechanisms involved in the fractionation of hydrogen isotopes between water and lipid are not fully resolved. We provide data aimed at understanding this fractionation in the Gram-positive obligately thermophilic anaerobe, *Thermoanaerobacterium saccharolyticum*, by comparing a wild-type strain to a deletion mutant in which the *nfnAB* genes encoding electron-bifurcating transhydrogenase have been removed. The wild-type strain showed faster growth rates and larger overall fractionation 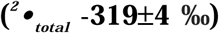 than the mutant strain 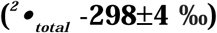. The overall trend in growth rate and fractionation, along with the isotopic ordering of individual lipids, is consistent with results reported for the Gram-negative sulfate reducer, *Desulfovibrio alaskensis* G20.

## 1. Introduction

The fractionation of hydrogen isotopes between environmental water and microbial biomass lipids correlates with central energy metabolism in many aerobic and some anaerobic bacteria (Dawson et al., 2015; Heinzelmann et al., 2015; Osburn et al., 2016; Zhang et al., 2009). The correlation has been inferred to relate to the mechanisms controlling the production of intracellular electron carriers such as NADPH and NADH. In some anaerobic bacteria the pattern of fractionation is more complicated, and does not strongly correlate with central carbon metabolism (Dawson et al., 2015; Leavitt et al., 2016; Osburn et al., 2016). One potential explanation for this complexity relates to the importance of flavin-based electron bifurcation by transhydrogenase in anaerobes (Demmer et al., 2015). These enzymes may impose a large isotope effect, which could overprint signals that relate primarily to carbon metabolism. Examination of transhydrogenase mutants in *Desulfovibrio alaskensis* G20 showed that on substrates such as malate and fumarate, perturbed transhydrogenase significantly affected the •^2^H values of lipids (Leavitt et al., 2016).

A more complete understanding of factors that impact 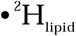 might be achieved by examination of microbial strains with different strategies for NAD(P)H regulation. The production of NADPH is critical for lipid biosynthesis. This cellular metabolite can derive from multiple sources, including reactions of central carbon metabolism, production from NADH via transhydrogenase, and production from NADH by NAD kinases. Three types of NAD kinases have been described (Kawai and Murata, 2008), with subcategories found in (i) Gram positive (+) bacteria and archaea, (ii) eukaryotes, and (iii) Gram negative (−) bacteria. In Gram(+) bacteria such as *Thermoanaerobacterium saccharolyticum*, NAD kinase can use ATP or polyphosphate as a P source. Few data exist on hydrogen isotopic fractionation in Gram(+) bacteria (Valentine et al., 2004). In this study, we apply a molecular genetic approach to examine hydrogen isotopic fractionation in a model Gram(+) organism, *T. saccharolyticum*. We compare the wild-type strain to a transhydrogenase-deficient mutant to determine phenotypic effects on growth rate, lipid profile, and magnitude of hydrogen isotopic fractionation between medium water and lipid. Our findings show patterns similar to those observed for *D. alaskensis* G20 (Leavitt et al., 2016).

## 2. Methods

*T. saccharolyticum* strain JW/SL-YS485 (wild type) was cultivated in parallel with a recently reported NfnAB transhydrogenase-deficient mutant (Lo et al., 2015), strain LL1144 (*•nfnAB*::Kanr). Triplicate cultures of each were grown at 55 °C in 150 ml glass bottles with a 50 ml working volume in MTC defined medium on 5 g/l cellobiose, as detailed in the Supplement. Cells were harvested at early stationary phase by way of centrifugation and were lyophilized. Lipid extraction, derivatization and analysis protocols were identical to those reported by Leavitt et al. (2016). Lipid retention times and peak areas were determined by gas chromatography-mass spectrometry (GC-MS), the •^2^H values of lipids measured by GC isotope ratio mass spectrometry (GC-IRMS) and the •^2^H of water samples by duel-inlet IRMS and cavity ring-down spectroscopy (CRDS), following Leavitt et al. (2016). The •^2^H values are reported relative to V-SMOW (Vienna Standard Mean Ocean Water) and fractionation is reported as apparent fractionation between medium water and lipid from the equation: 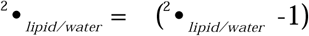, where 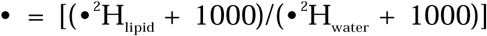. Each lipid from each culture sample (representing each individual biological triplicate) was measured 14 to 20 times.

## 3. Results

The doubling time of the wild-type strain was 0.33 ± 0.10 h^−1^, compared vs. a slower growth rate of 0.10 ± 0.01 h^−1^ for the *•nfnAB* strain (Fig. 1). The wild-type strain demonstrated a longer lag phase, perhaps because it was inoculated at a lower initial cell density than the mutant. The maximum optical density (OD) for the wild-type was nearly 3-times that of the mutant, with average final OD_600_: wild-type = 1.04 (±0.03) vs. *•nfnAB* = 0.37 (±0.01), representing biological triplicates of each strain (Fig. 1).

**Fig. 1.**
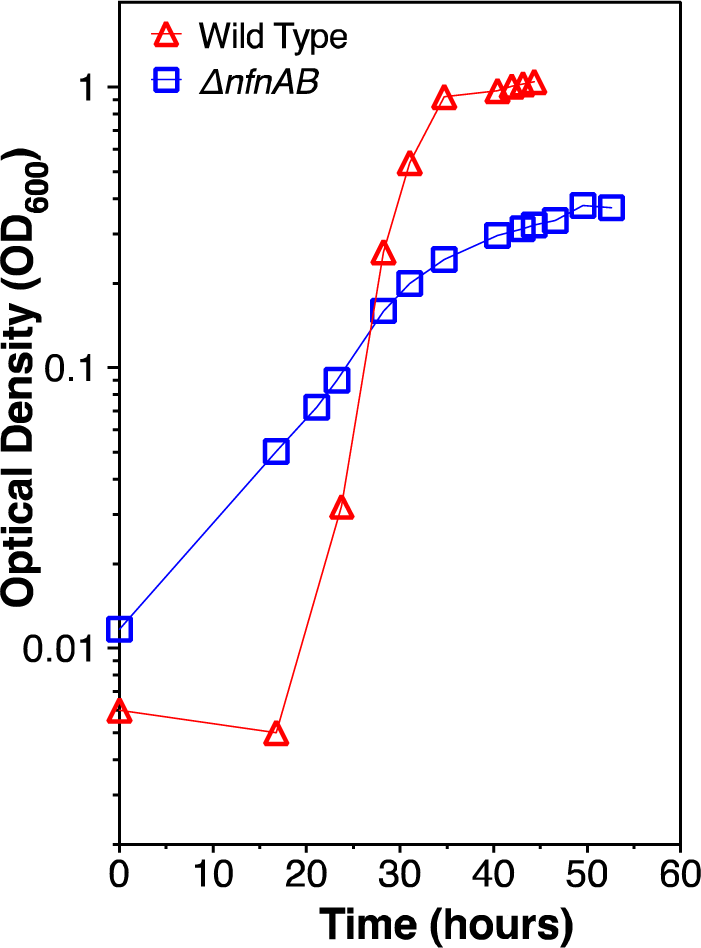
Growth curves and calculated doubling times for wild-type and mutant (avg. of triplicate growth experiments).

The lipid profile of *T. saccharolyticum* was similar to what has been previously reported from this genus (Jung and Zeikus, 1994). The strain produced abundant *n*-C_16_ fatty acids (FA) along with branched *iso-* and *anteiso-* C_15_ and C_17_ FA. Smaller amounts of *n*-C_14_ FA were detected, along with a long-chain dicarboxylic acid. The mass spectrum of the latter was consistent with one reported from *T. ethanolicus* (Jung and Zeikus, 1994). The wild-type had elevated concentrations of the *iso-* FA relative to the mutant, but the lipid profiles were otherwise similar (Fig. 2).

**Fig. 2.**
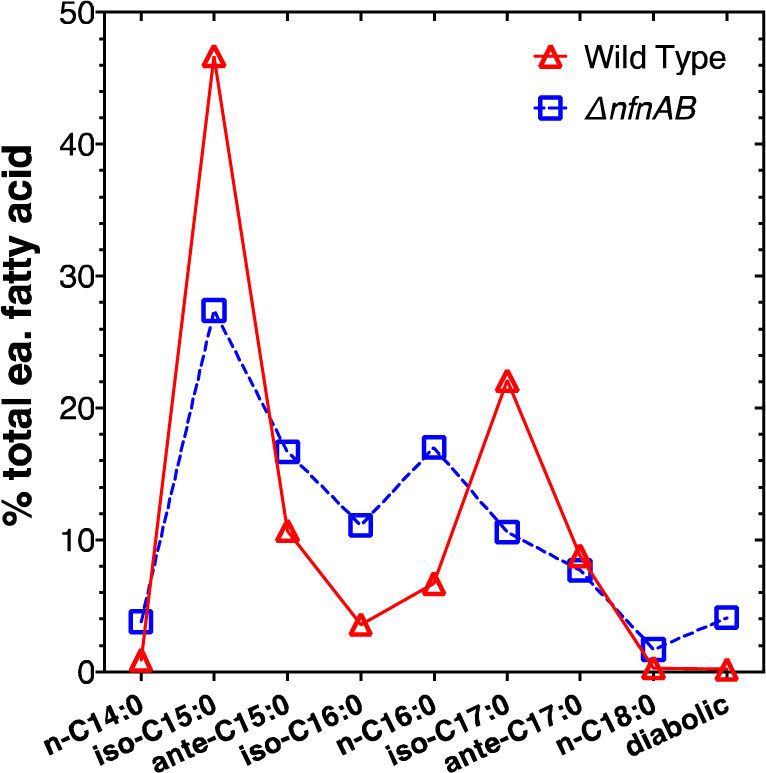
Lipid abundance profiles for wild type and mutant (avg. of triplicate growth experiments).

The mass-weighted average hydrogen isotopic fractionation between water and lipid 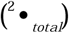 was greater for the wild type (−319±4 ‰) than for *•nfnAB* (−298±4 ‰). The fractionation (^2^•) for each individual lipid was also greater in the wild type than in the mutant (Fig. 3). The isotopic ordering of individual lipids 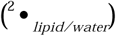 was similar for both strains, with *anteiso*- lipids depleted relative to *iso*-and straight chain FA. The relative ordering from most depleted lipid (*anteiso*-C_15:0_) to most enriched (iso-C_17:0_), was nearly identical for both wild-type and mutant (Fig. 3).

**Fig. 3.**
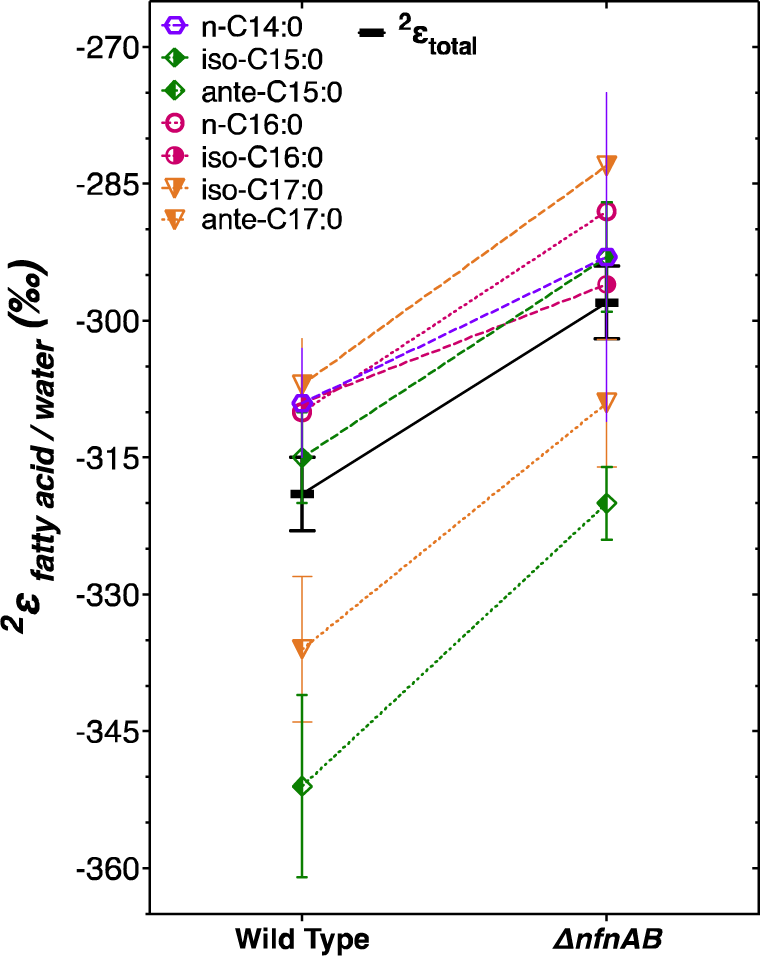
H isotope fractionations between FA and water. Black horizontal bar, weighted mean for each strain. Vertical bars, standard mean error (SME) for all biological (N = 3) and technical replicates (n = 14 to 20).

**Fig. 4.**
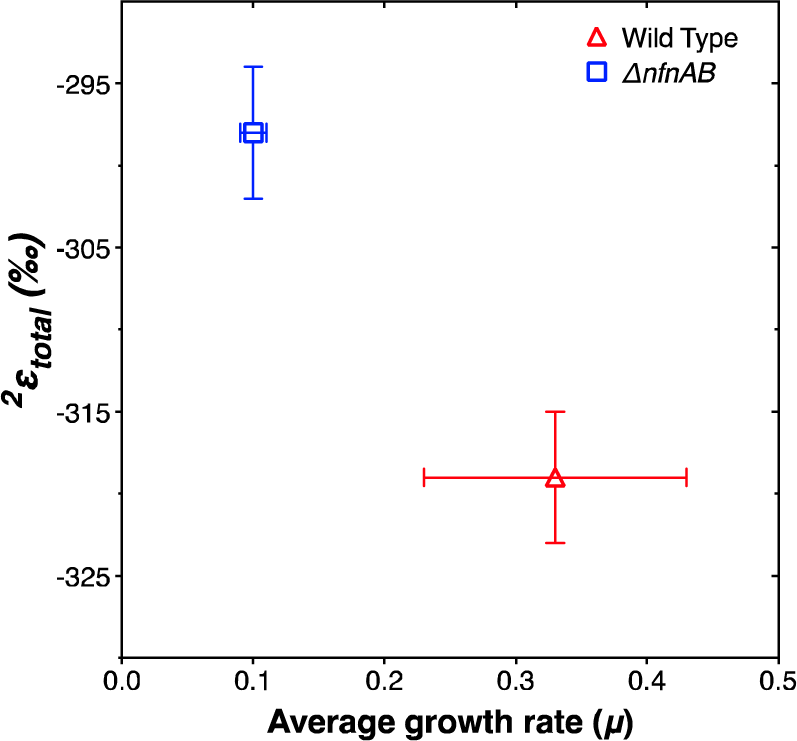
Weighted H-isotopic fractionation between FA and water for each strain vs. average doubling time.

## 4. Discussion

Observation of the 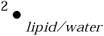 and 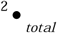 in wild-type and *nfnAB* transhydrogenase mutant strains of *D. alaskensis* G20 revealed that faster growing strains were more depleted in ^2^H than the slower strains (Leavitt et al., 2016). *T. saccharolyticum* also showed similar relationships between growth rate and fractionation. Whether this pattern can be attributed to a similar role for the influence of transhydrogenase on 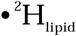, a consistent relationship with growth rate and 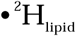, or a more nuanced relationship due to changes in both NfnAB activity and growth rate, remains unresolved. Deconvoluting these possibilities will require steady-state experiments with both strains cultured in parallel at a fixed growth rate. Growth rate effects have been observed on 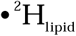 in haptophyte algae (Sachs and Kawka, 2015; Schouten et al., 2006), and chemostat experiments have been used to understand fractionation as a function of rate in other isotope systems (Leavitt et al. 2013).

Another commonality between *T. saccharolyticum* and *D. alaskensis* is the ^2^H depletion in the *anteiso*-FA relative to the other FA (Fig. 3). Leavitt et al. (2016) suggested that this depletion might originate in the biosynthesis of *anteiso*-FA from 2-methylbutyryl-CoA derived from isoleucine. This explanation could also be invoked for *T. saccharolyticum*, and compound-specific •^2^H measurements of amino acids might provide valuable constraints on the isotopic ordering among lipids. A recent study of the H isotopic compositions of individual amino acids in *Escherichia coli* grown on glucose and tryptone showed that isoleucine was depleted in ^2^H relative to leucine by ca. 100‰ (Fogel et al., 2016).

## 5. Conclusions

Deletion of the electron-bifurcating transhydrogenase, NfnAB, slows growth rate and decreases the magnitude of 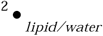 and 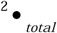 when *T. saccharolyticum* is grown on a defined medium in batch culture. The relative ordering of 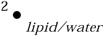 is similar in both strains. These patterns of fractionation and isotopic ordering are similar to recent observations of the heterotrophic sulfate reducer *D. alaskensis* G20. The consistency of results across these taxa supports a role for NfnAB in determining the H isotopic composition of lipids in obligate anaerobes. However, to better constrain these observations, and isolate the effect of growth rate, continuous culture (chemostat) experiments are necessary. Similar work with a broader array of transhydrogenase-containing microbes would be helpful, including organisms utilizing families of transhydrogenases other than NfnAB-class. Such experiments can place further constrains on the mechanism(s) of lipid H-isotopic fractionation.

## 6. Supplementary information

All supplemental methods and data are archived at: 10.6084/m9.figshare.4598224.

## Acknowledgments

We thank K. Grice and J. Maxwell for editorial handling and an anonymous reviewer whose comments helped improve the manuscript. We are grateful to M. Seuss (Washington University) for assistance with lipid analysis, X. Feng (Dartmouth College) and M. Osburn (Northwestern University) for water H-isotope analysis and the Lynd lab (Dartmouth College) for assistance with growth experiments. Funding was provided by U.S. NASA Exobiology grant 13-EXO13-0082 (A.S.B., W.D.L.), and the Fossett Postdoctoral Fellowship from Washington University in St. Louis (W.D.L.). The BioEnergy Science Center is supported by the Office of Biological and Environmental Research in the U.S. Department of Energy Office of Science. The manuscript has been authored in part by Dartmouth College under Contract no. DE-AC05-00OR22725 with the U.S. Department of Energy.

